# Robust T-cell signaling triggered on soft polydimethylsiloxane-supported lipid-bilayers

**DOI:** 10.1101/2020.06.05.134684

**Authors:** Anna H. Lippert, Ivan B. Dimov, Alexander Winkel, James McColl, Jane Humphrey, Kevin Y. Chen, Ana Mafalda Santos, Edward Jenkins, Kristian Franze, Simon J. Davis, David Klenerman

## Abstract

The T-cell receptor (TCR) is thought to be triggered either by mechano-transduction or local tyrosine phosphatase exclusion at cell-cell contacts. However, the effects of the mechanical properties of activating surfaces have only been tested for late-stage T-cell activation, and phosphatase segregation has mostly been studied on glass-supported lipid bilayers that favor imaging but are orders-of-magnitude stiffer than typical cells. We developed a method for attaching lipid bilayers to polydimethylsiloxane polymer supports, producing ‘soft bilayers’ with physiological levels of mechanical resistance (Young’s modulus of 4 kPa). Comparisons of T-cell behavior on soft and glass-supported bilayers revealed that early calcium signaling is unaffected by substrate rigidity, implying that early steps in TCR triggering are not mechanosensitive. Robust phosphatase exclusion was observed on the soft bilayers, however, suggesting it likely occurs at cell-cell contacts. This work sets the stage for an imaging-based exploration of receptor signaling under conditions closely mimicking physiological cell-cell contact.

## Introduction

Cells and tissues exhibit a wide range of stiffnesses from 100 Pa for lymphocytes, up to GPa for bone^1^. This suggests that cells may have to be mechanosensitive to correctly navigate and/or respond to their local environments. It could be expected that lymphocytes would comprise a special case of mechanosensitive cells, since they traverse great distances across multiple organs and tissues in the course of interrogating other cells for signs of infection and malaise.

Accordingly, multiple groups have shown that lymphocytes, i.e. both T cells ^2,3^ and B cells ^4,5^ are affected by surface mechanical properties. For example, T cells interacting with elastomer micropillar arrays undergo cytoskeletal changes and intracellular signaling responses that vary with pillar length and flexibility ^6^. Substrate stiffness also modulates T-cell migration and morphology and alters the expression of T-cell receptor (TCR)-induced immune system-, metabolism- and cell cycle-related genes ^7,3^. But it is also known that antigen-presenting cells (APCs) change their stiffness in the course of the inflammatory response ^1,7^, and that cancer cells can be up to five-fold softer than their healthy counterparts ^8,9^. Therefore, T cells may have to accommodate significant variation in surface rigidity in the course of performing their essential protective functions. How T cells would parse these conflicting requirements is unknown.

Consistent with the reported sensitivity of T cells to substrate rigidity, a considerable body of data implies that T cells use force-sensing mechanisms acting through the TCR to initiate signaling ^10–13^. Paradoxically, however, T-cells are unusually soft, implying that membrane deformability is important. A second explanation for signaling that would exploit this proposes that the TCR relies on the passive exclusion of large receptor-type tyrosine phosphatases, such as CD45, from small regions of contact with APCs where the TCR engages its ligands, driven only by bond formation between TCR-sized adhesion proteins such as CD2 and its ligands ^14,15^. Local removal of the phosphatases is suggested to favor receptor phosphorylation by kinases attached to the inner leaflet of the T-cell membrane, with contacts needing to be small to ensure responses are antigen-dependent ^16,17^. CD45 is readily seen to exit from μm-sized regions of T-cell contact with glass surfaces and bilayers ^16–18^, but when live T-cell/APC contacts were imaged using lattice light-sheets, CD45 exclusion was not observed ^19^. Explanations for this are that cell surfaces are too soft to effect phosphatase exclusion, or that it is too short-lived or occurs on too small a length-scale to be imaged using light-sheets.

Imaging cells on model surfaces comprising glass-supported lipid-bilayers has previously offered the best opportunity to study the early stages of T-cell activation at high spatial and temporal resolution, but these surfaces are orders-of-magnitude stiffer than typical cell surfaces. Here, we establish a new method for attaching bilayers to soft polydimethylsiloxane (PDMS) supports. We use the soft bilayers to probe the stiffness-dependence of surface reorganization and receptor-proximal signaling by T cells.

## Results

### Creation and characterization of PDMS-supported lipid bilayers

To study T-cell signaling on surfaces that had softnesses comparable to those of lymphocytes but retained the advantages of supported lipid bilayers (SLBs) for imaging, we formed SLBs on PDMS supports. Whereas previous methods of attaching SLBs to PDMS were reliant on solvent or plasma treatments that influence the stiffness of the PDMS support ^27^, our method is both simple and non-perturbative. The method involves an overnight incubation of PDMS-presenting slides with a 1 mM calcium chloride solution prior to bilayer formation using small unilamellar vesicles (Fig. 1a). The bilayers thus formed can then be functionalized using histidine-tagged proteins in the manner of glass-supported bilayers. We found that lipid bilayers could be formed on PDMS supports of different stiffness, ranging from 4 kPa to 1 MPa. Atomic force microscopy (AFM) measurements (Suppl. Fig. 1a) confirmed that the PDMS gels exhibited predominantly elastic responses with minimal creep (Suppl. Fig. 1b), and that calcium treatment did not lead to an increase in gel stiffness (Suppl. Fig. 1c,d). Typically, the PDMS supports for the bilayers were ~50 μm in depth. For most of our functional experiments comparing the glass-supported and soft bilayers, PDMS gels of 4 kPa stiffness were used, which matches the surface stiffness of mature dendritic cells (3.5 kPa) ^28^, the most relevant surface for understanding T-cell activation.

**Fig. 1:**
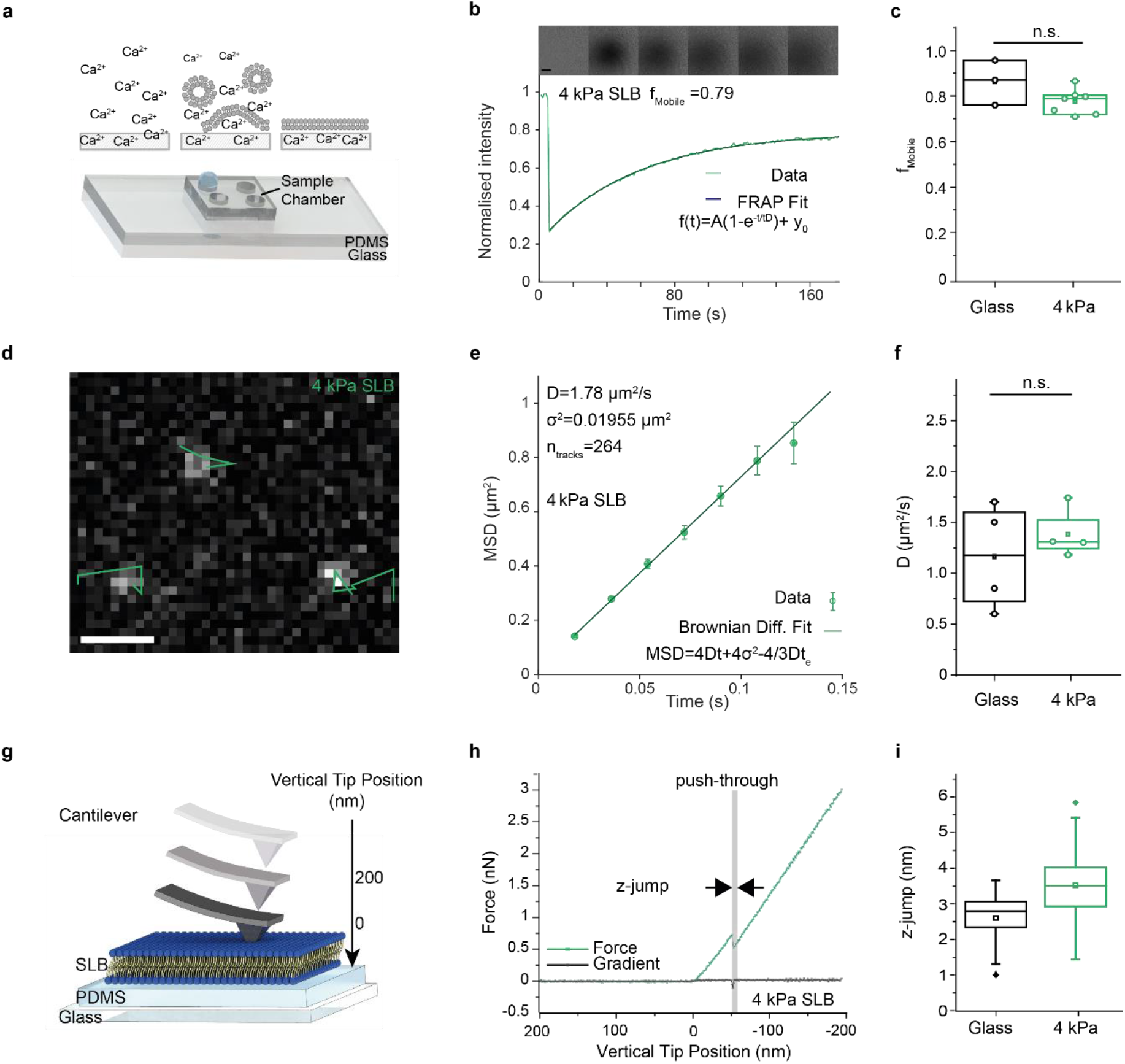
Formation and characterisation of PDMS-supported bilayers. **a**, Schematic representation of the experimental setup and the formation process. **b** Representative FRAP curve and fit from a SLB on 4 kPa PDMS gel with raw data showing the fluorescence recovery after photobleaching, above. Scale bar 10 μm. **c** Mobile fraction *f*_mobile_ assessed using FRAP. Each data point represents one independent experiment, performed at room temperature. **d-f** Single-molecule diffusion measurement of lipids in the PDMS supported bilayers. **d** Representative TIRF image and tracks (green) of single oregon-labelled lipids diffusing in a bilayer formed on a 4 kPa PDMS gel. Scale bar 1 μm. **e** Representative MSD plot and fit of data acquired on a SLB formed on 4 kPa PDMS gel (264 tracks). **f** Comparison of the diffusion coefficients of lipids on SLBs supported by glass and 4 kPa PDMS gels. Each data point represents one independent measurement of lipid diffusion at RT (n_tracks_ > 100). **g-i** Mechanical characterisation of SLBs on PDMS *via* AFM. **g** Schematic representing the AFM cantilever approaching and contacting the SLB and then the PDMS. **h** Representative force curve (green) on a 4 kPa PDMS gel. Push-through events are detected via changes in the gradient of the force curve (black). **i** Boxplots of SLB thickness (z-jump). Boxes indicate the 25% and 75% quartile, the horizontal line the mean, and whiskers the 1.5 IQR (number of force curves analysed: 52 (4 kPa PDMS), 30 (glass), from 3 independent experiments). The Mann-Whitney U test was used for statistical comparisons.

To confirm that the bilayers were fluid and mobile, the diffusion coefficient of the lipids along with their mobility were assessed using FRAP (Fig. 1b,c; Suppl. Fig. 2a) and single-molecule tracking (Fig. 1d-f; Suppl. Fig. 2b,c; Suppl. Video 1). We found that 80-90% of the lipids in the bilayers on both 4 kPa and 1 MPa PDMS supports were mobile, with diffusion coefficients of 1.5 μm^2^/s (Fig. 1e,f). These values were similar to those for bilayers formed on glass (Fig. 1c,f; Suppl. Fig. 2a,c). To establish that a single bilayer and not multiple layers had formed on the PDMS supports we performed AFM force curve measurements, analysing single push-through events using a pyramidal AFM cantilever (Fig. 1g). In these experiments the cantilever was first brought into contact with the bilayer and then, with increasing force, pushed through the bilayer and into contact with the underlying PDMS support. On 4 kPa PDMS-supported bilayers, 51 of 52 push-through experiments produced a single push-through event (see, *e.g.*, Fig. 1h), with just one force curve showing two sequential push-through events. This indicated the formation of single bilayer sheets on the PDMS, instead of multiple bilayers. Our glass-supported bilayers also behaved as expected (Suppl. Fig. 3a). Both 4 kPa PDMS- and glass-supported bilayers exhibited similar resistance to push-through (Suppl. Fig. 3b) and had similar depths (z jumps; Fig. 1i) that were in good agreement with the reported values of ~3.9 nm for POPC bilayers ^29,30^. The ratio of the slopes before and after push-through for the 4 kPa PDMS-supported bilayers (Fig. 1h; Suppl. Fig. 3c) was close to 1, indicating that the bilayer and underlying support were of similar stiffness.

### Properties of contacts formed by T cells interacting with PDMS-supported bilayers

One of the advantages of PDMS as a supporting substrate is that it has a refractive index similar to glass, allowing TIRF imaging. Using this type of imaging, therefore, we set out to determine the extent to which substrate stiffness affects the nature of the contacts Jurkat T-cells form with model surfaces.

For this we imaged the behaviour of the large glycocalyx component/receptor-type tyrosine phosphatase, CD45, expressed by the T cells, and the small, histidine-tagged adhesion protein rat (r) cluster of differentiation antigen 2 (rCD2), inserted into the SLBs to allow for the formation of adhesive contacts (Fig. 2a). CD45 was indirectly labelled with the Fab fragment of the Gap8.3 antibody ^31^, whereas CD2 was labelled directly on lysines with Alexa-647 dye.

**Fig. 2:**
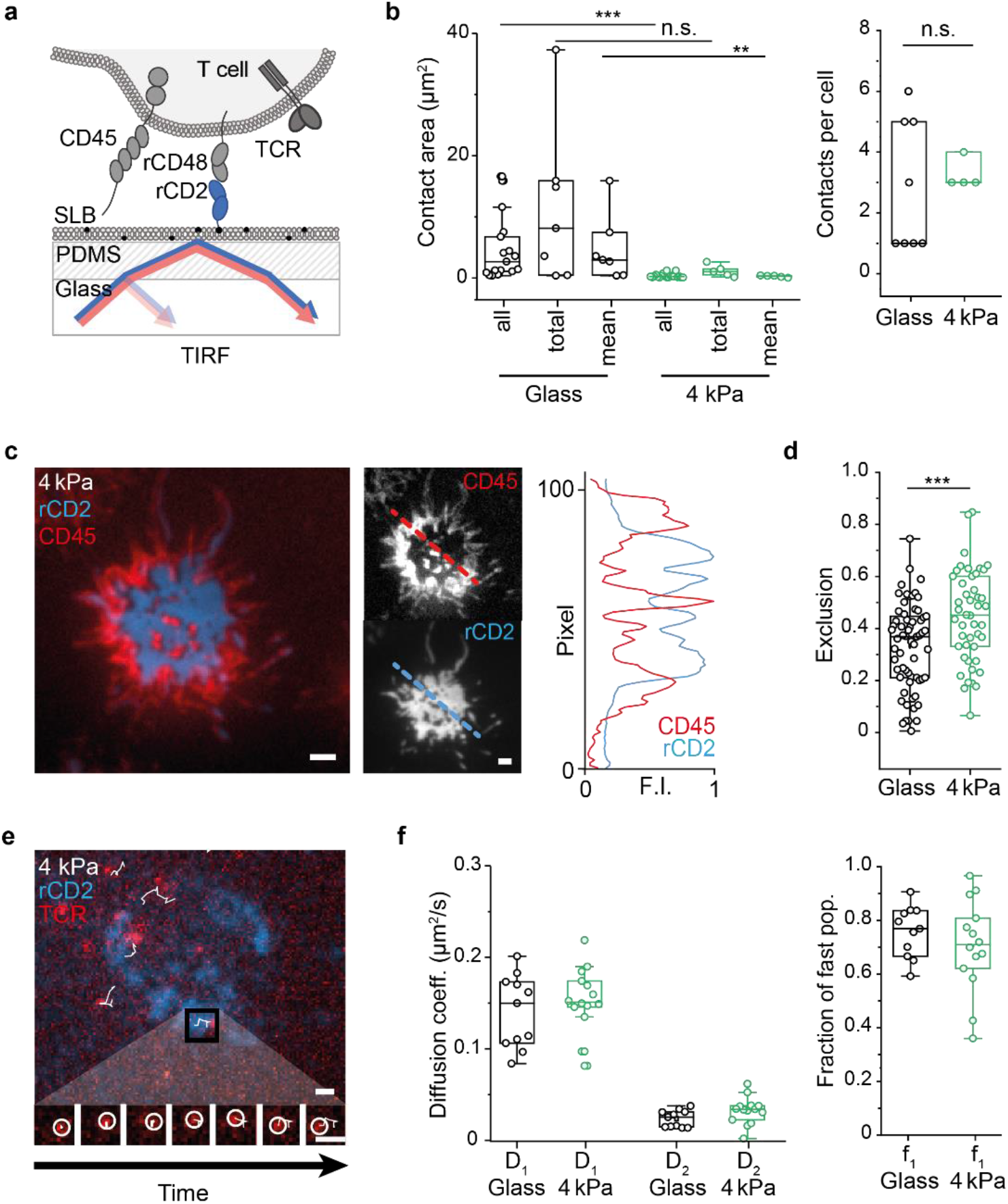
Properties of contacts with the PDMS-supported bilayers presenting the adhesion protein rCD2. **a** Schematic of the experiment. Cells expressing signaling-disabled rCD48 were deposited onto SLBs presenting rCD2. Cells were labelled with Gap8.3 anti-CD45 Fab or UCHT1 anti-CD3 Ab. **b** Boxplots comparing rCD2 contacts one frame prior to calcium peak. Areas for all individual contacts (all), total contact area per cell (total), and average contact size (mean) were each measured. **c** Numbers of contacts per cell. For **b** and **c**, n_cells_=7 (glass) and n_cells_=5 (PDMS) acquired during 3 independent experiments. Experiments were performed at 37°C. **d** Overlay and individual TIRF images showing CD45 (red) and rCD2 (blue) fluorescence with a line profile for normalised pixel intensity (dashed lines). Scale bars are 1 μm. **e** Boxplots of CD45 exclusion. Boxes indicate the 25% and 75% quartile, the horizontal line the mean, and whiskers the 1.5 IQR. Each data point corresponds to one cell; n_cells_=63 (glass) and n_cells_=47 (PDMS) acquired from 3 independent experiments at RT. **f** Tracking of TCRs labelled with UCHT1 anti-CD3∊ Fab tagged with Alexa488 on a 4 kPa PDMS bilayer. Shown is a z-projection of the acquired data (red) with TCR tracks overlayed (white) and the rCD2 contact (blue). Shown below is an example of a single TCR diffusing over time (interval 31 ms). Scale bars are 1 μm. **g** Cell-wise 2-component JD fit of the JD distributions of both glass-supported bilayer tracking data (n_cells_=12, n_tracks_ per cell>30, mean track length over all cells: 20.8±5.4, mean SNR: 7.8±1.4) and 4 kPa PDMS-supported bilayer tracking data (n_cells_=15, n_tracks_ per cell>30, mean track length over all cells: 15.9±4.2, mean SNR: 6.0±0.9) obtained in 2 independent experiments performed at 37°C. Shown are D1, D2 and f1. The Mann-Whitney U test was used for statistical analysis with *:<0.05, **:<0.01, ***:<0.001.

We imaged Jurkat T-cells expressing rCD48, the ligand of rCD2, as they formed contacts with rCD2-functionalized bilayers formed on glass and on 4 kPa PDMS gels. T cells readily formed single or multi-focal contacts on both the soft and stiff bilayers. However, significantly smaller contacts, in terms of both the total area of cellular contact and the size of individual contacts, formed in the case of cells interacting with the 4 kPa PDMS-supported bilayers versus the glass-supported SLBs, both ‘early’ (at the time of calcium signaling; Fig. 2b) and ‘late’ (10 minutes after contact with the bilayers; Suppl. Fig. 4a). However, the numbers of contacts formed per cell were not significantly affected by bilayer stiffness (Fig. 2c; Suppl. Fig. 4a). Cell contacts with the bilayers were marked by the accumulation of fluorescent CD2: an example of an early contact formed on a 4 kPa PDMS-supported bilayer is shown in Fig. 2d. Notably, CD45 was excluded from regions of CD2 accumulation (Fig. 2d). Time-lapse imaging (Suppl. Fig. 5a) was used to follow the formation of contacts, and this showed that CD45 was excluded very early (Suppl. Fig. 5b). Surprisingly, the extent of CD45 exclusion was slightly greater on soft versus stiff bilayers (Fig. 2e). Single-molecule tracking using two-colour TIRF imaging of single TCRs within the rCD2-accumulating contacts revealed that receptor diffusion was rapid (0.15 μm^2^/s), as reported previously ^32^, and unaffected by bilayer stiffness (Fig. 2f,g; Suppl. Fig. 4b; Suppl. Video 2). Similarly, large fractions of the TCRs were mobile in both types of bilayers.

### Signaling on PDMS-supported bilayers

We next sought to determine the influence of substrate stiffness on T-cell activation. Whereas previous studies examined the stiffness-dependence of activation using late readouts such as IL-2 (refs ^3,33^), we focussed on early T-cell activation, using calcium signaling responses as a proximal readout of receptor triggering (Fig. 3a). We used T cells expressing the 1G4 TCR, which binds the NY-ESO 9V peptide ^34^ presented by the HLA-A2 major histocompatibility complex (pMHC). To detect calcium signaling we used cells expressing the high signal-to-noise green fluorescent protein-based calcium reporter, GCAMP ^21^ (for cells responding to pMHC), or cells labelled with the calcium reporter Fluo-4 (all other experiments).

**Fig. 3:**
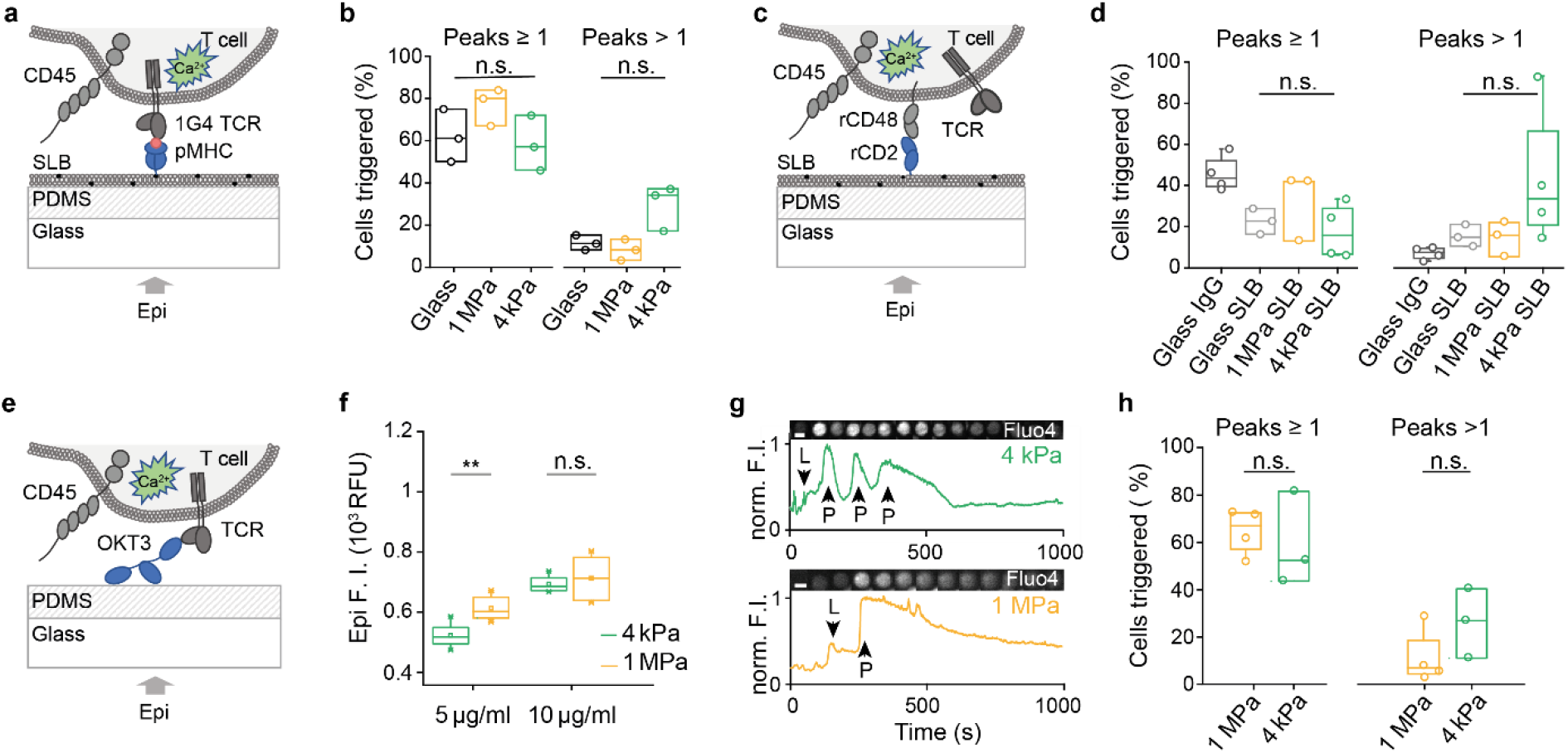
T-cell calcium signaling is largely independent of substrate stiffness. **a-b** Ligand-dependent calcium signaling on pMHC coated bilayers. **a** Schematic representation of the experiment. T cells expressing the genetically encoded calcium sensor GCaMP and the 1G4 TCR complex were imaged on pMHC-coated bilayers. **b** Boxplots show the fraction of cells exhibiting a calcium response (peaks ≥ 1) and the fraction of triggering cells exhibiting multiple peaks (peaks >1), for cells interacting with glass-supported and 1 MPa and 4 kPa PDMS-supported bilayers. **c-d** Ligand-independent calcium signaling. **c** Schematic of the experiment. Fluo-4 calcium reporter loaded Jurkat T-cells expressing signaling-disabled rCD48 were imaged on rCD2-presenting SLBs. **d** Boxplots showing fractions of cells exhibiting a calcium response (peaks ≥ 1) and those exhibiting multiple peaks (peaks > 1) on glass coated only with unspecific antibody (bovine IgG), and on bilayers supported by glass or 1 MPa and 4 kPa PDMS. Each data point represents an independent experiment with n_cells_>50. Boxes indicate the 25% and 75% quartile, the horizontal line the mean, and whiskers the 1.5 IQR. **e-h** Calcium responses of T cells on PDMS coated with activating OKT3 anti-CD3∊ antibody. **e** Schematic of the experiment. T cells were preloaded with Fluo-4 prior to imaging their calcium responses on OKT3-coated PDMS. **f** PDMS gel stiffness did not influence antibody loading at the concentration used (10 μg/ml). Gels were incubated with fluorescently labelled antibody overnight, washed and the fluorescence intensity measured using epi-illumination (2 gels per condition, 4 fields of view). **g** Example calcium signaling traces for cells contacting OKT3 antibody-coated 4 kPa and 1 MPa PDMS gels; L, cell landing, P, calcium peaks. Fluo4 fluorescence is shown above the trace. Scale bar 10 μm. **h** Fraction of cells exhibiting a calcium response (peak ≥ 1) and fraction of responding cells that displayed multiple peaks (peaks > 1). Each data point in the boxplot represents a single experiment with n_cells_>50 for each point. All experiments were performed at 37°C. The Mann-Whitney U test was used for statistical analysis with *:<0.05, **:<0.01.

T cells were first imaged as they contacted bilayers presenting pMHC. On glass- and on 4 kPa and 1 MPa PDMS-supported bilayers, similar large fractions of cells (60-80%) produced calcium signaling responses (Fig. 3b). However, we also noticed that a relatively small fraction of the cells exhibited blinking behaviour, rather than single sustained responses (Fig. 3b). The fraction of cells exhibiting the blinking behaviour was higher on the 4 kPa PDMS-supported bilayers than for 1 MPa PDMS (~30% versus ~10%), but the trend did not reach statistical significance. On bilayers presenting histidine-tagged rCD2 (Fig. 3c), we saw weaker but non-trivial levels of TCR-ligand independent signaling, as reported previously (Fig. 3d; Suppl. Videos 3,4) ^16,35^. A time-course analysis showed that even very small contacts on 4 kPa bilayers were capable of initiating ligand-independent calcium signaling (Suppl. Fig. 6). These types of responses, which were initially observed by Chang *et al.* ^16^ for cells interacting with glass surfaces coated with unspecific (bovine) IgG (see also Fig. 3d) and were attributed to the effects of local CD45 exclusion, exhibited the same tendency toward increased blinking behaviour as substrate stiffness decreased. Other parameters, such as calcium peak height, time to trigger and integrated peak intensity were similar for the glass- and 4 kPa PDMS-supported bilayers (see Suppl. Fig. 7). Finally, similar signaling behaviour was observed on PDMS surfaces of differing stiffness derivatised with the widely used, TCR triggering anti-CD3ε antibody, OKT3 (Fig. 3e-h), indicating that all routes to TCR triggering are indifferent to surface rigidity.

## Discussion

Although at times controversial it now appears that, like other receptors such as cytokine receptors and GPCRs, individual TCRs initiate downstream signaling after binding their ligands ^36^. But unlike these other receptors, the ligands of the TCR are exclusively cell-bound. To understand how TCRs are triggered, therefore, it will be necessary to observe events at the interface of interacting cells, at the single-molecule level. For now, the only way to study cell surface phenomena at the single-molecule level is to use TIRF-based imaging, and model cell surfaces. TIRF imaging of glass-supported lipid bilayers used as model APC surfaces, a method pioneered by Dustin and collaborators ^37^, has transformed our understanding of T-cell activation ^38^. A special advantage of the bilayer approach is that the bilayers can be easily and systematically functionalized by attaching the soluble extracellular regions of receptor ligands and other components of the membrane, allowing quantitative analysis of T-cell responses. In recent years, however, it has become apparent that T-cell behavior is affected by the mechanical properties of surfaces they encounter ^3,33,39,40^ and glass is, of course, orders-of-magnitude stiffer than all eukaryotic cells.

We sought better mimics of the cell surface that preserve the utility of bilayer technology. We show here that bilayers can be attached to PDMS gels without significantly affecting their stiffnesses. We generated “soft” bilayers differing in stiffness across 2-3 orders of magnitude, including supports closely matching the stiffness of mature dendritic cells, the archetypal antigen-presenting cell (3-4 kPa) ^28^. A key advantage of PDMS substrates is that they allow TIRF imaging. Single-molecule tracking and diffusional analysis of labelled lipids in the bilayers allowed us to confirm that the surfaces were fluid, and AFM push-through experiments showed that we had produced single bilayers of the expected height. All measured parameters agreed well with the published literature ^29,30^, indicating that with respect to, *e.g.*, thickness of the lipid bilayer and the diffusion of lipids, the biophysical properties of our soft PDMS-supported bilayers matched those on glass supports.

Illustrating their utility, we used the new bilayers to probe the stiffness-dependence of surface reorganization and receptor-proximal signaling by T cells, by comparing T-cell behavior on glass- and 4 kPa PDMA-supported bilayers. TCR diffusion in the absence of ligands was the same on both bilayers, and we could readily observe the patterns of μm-scale spatial reorganisation of key surface proteins on soft bilayers that had been reported previously, including the local accumulation of a small adhesion protein, rCD2, and the exclusion from cellular contacts of the large glycocalyx element, CD45 ^16,41–43^. Whereas we had previously observed the local segregation of CD45 from regions of contact of T cells with glass surfaces and supported bilayers ^16^, Cai *et al.* did not observe CD45 exclusion from contacts made by T cells interacting with APCs, imaged using lattice light-sheets ^19^. Whatever the reasons are for this, it seems unlikely that differences in the stiffness of cells and glass surfaces are the explanation because, at stiffness levels closely matching those of APCs, we saw robust CD45 exclusion.

Previous work also showed that substrate stiffness and T-cell activation, measured as IFNγ and TNFα production, and CD25 expression across time-scales of days, were correlated ^7^. In contrast, our measurements of intracellular Ca^2+^ increases, which reports TCR-proximal signaling ^44,45^, indicated that the earliest stages of T-cell signaling are indifferent to substrate stiffness. There was no reduction in the number of cells responding on 4 kPa and 1 MPa PDMS-supported bilayers versus glass-supported bilayers, and only a modest trend towards an increase in fluctuating Ca^2+^ responses, which are thought to correspond to weaker or incomplete T-cell activation ^46,47^. We obtained similar results for signaling initiated by an immobilised antibody, and for triggering induced in the absence of ligands. Overall, it seems unlikely that differences in TCR triggering are responsible for the late-acting effects of substrate stiffness identified by Saitakis *et al.*, the source of which is presently a mystery.

These results suggest that the TCR is not sensitive to surface rigidity in the manner typical of mechanotransducers. This might crucially allow T cells to respond to APCs or tumours whose surface stiffnesses can vary over a substantial range ^1,7–9^. It could be argued that 4 kPA is stiff enough to create the tension needed to trigger the TCR, but even lower levels of rigidity have been shown previously to allow TCR triggering. For example, the passive contact of T cells with small TCR ligand-presenting unilamellar vesicles have been shown to initiate spontaneous CD45 exclusion and receptor triggering, with similar findings obtained for receptor triggering in B cells and mast cells ^48^. It also bears repeating that on glass-supported bilayers, TCRs insulated from forces by adhesion molecules also inducing the exclusion of CD45, are spontaneously triggered wholly in the absence of ligands ^16^. With the advent of PDMS-supported soft bilayers, the stage is now set for the systematic, imaging-based exploration of the receptor triggering problem, under conditions closely mimicking physiological cell-cell contact.

## Methods

### PDMS Fabrication

For the fabrication of 4 kPa and 1 MPa gels, two commercially available polymer reagents were mixed as follows: Silgard184 (Sigma Aldrich) was mixed in a 1:10 ratio (curing agent: monomer) to prepare 1 MPa gels. NuSil Gel 8100 (NuSil) was prepared following the manufacturer’s instructions (1:1 A and B components). The gel mix was then supplemented with 1% of the prepared Silgard184 mix. After gentle stirring, the gels were spread on glass slides using razor blades, with 0.0625mm Tape (3M 810 TAPE Scotch^®^ Magic Tape) as spacer, and subsequently cured at 65°C for 13 h after which they were ready for experiments. The Young’s modulus of the soft PDMS gels was found to be 4.59± 0.35 kPa (n=16) and creep response experiments showed a low viscous contribution indicating that the gels were mostly elastic in their response (see Suppl. Fig. 1).

### PDMS stiffness measurements

All PDMS substrate stiffnesses were measured using either a JPK CellHesion 200 or JPK NanoWizard 3 AFM (Bruker) operated in Force Spectroscopy mode and Arrow TL1 cantilevers (NanoWorld) onto which a polystyrene bead (PS-R-37.0 - microParticles Gmbh) had been glued. PDMS stiffnesses were measured in PBS, with 250 μg/mL BSA to prevent tip-sample adhesion. Recorded indentations were processed using the JPK Data Processing software to extract the Young’s Modulus from the Hertz model ^20^. Force spectroscopy was also used to detect bilayer push-through events using SHOCON-10 cantilevers (AppNano). These were then analysed using a purpose-written script in Matlab.

### SUV preparation

Small Unilamellar Vesicle (SUV) solutions were prepared in glass vials where the amount of desired lipid was added from stock solutions using glass syringes. After drying the lipids under nitrogen flow and at least 1h inside a desiccator, PBS was added to the vial to yield 10 mg/ml SUV stock solutions. Following a short vortexing step, the solution was sonicated in a water bath until the solution became clear. Bilayers were prepared from SUV solutions containing POPC (1-palmitoyl-2-oleoyl-sn-glycero-3-phosphocholine, Avanti Polar Lipids, 850457C) and 0.01% Oregon Green DHPE (1,2-Dihexadecanoyl-sn-Glycero-3-Phosphoethanolamine, ThermoFisher, O12650) for the lipid diffusion experiments. To functionalize the bilayers with proteins, SUV solutions of POPC with DGS-NTA(Ni) (1,2-dioleoyl-sn-glycero-3-[(N-(5-amino1-carboxypentyl)iminodiacetic acid)succinyl] (nickel salt), Avanti Polar Lipids, 790404C) at 2%DGS-NTA(Ni), 98%POPC were prepared.

### Bilayer preparation on PDMS and glass

After overnight incubation of the PDMS slides with 1 mM CaCl_2_ solution, the slides were washed 3 times with PBS, and then SUV solution added at 1 mg/ml with additional calcium at 10 μM. After 30 min incubation and washing to remove free SUVs, proteins were added and incubated for 60 min. Here, proteins bind to the Ni^+^ chelating NTA lipid via 2xhexahistidine tags. Prior to experiments bilayers were washed 5 times with PBS to remove unbound proteins. Bilayers were either functionalized with 25 nM rCD2-Alexa647 (~100-200 molec/μm^2^) alone or with 70 nM pMHC (~600-800 molec/μm^2^). Bilayers were checked for mobility prior to each experiment. Bilayers on glass were prepared on piranha cleaned glass slides followed by a 30 min argon plasma treatment, before adding the SUV solution to the cleaned glass. After 30 min to allow bilayer formation on the glass, samples were washed 3 times with PBS.

### Cell culture

The Jurkat rCD48 cell line was generated *via* lentiviral transfections with the pHR lentiviral vector. The J8 Jurkat line was engineered to express the calcium indicator GCaMP7s^21,22^ for when needed, as well as the 1G4 TCR complex. All T cells were cultured in phenol red-free RPMI supplemented with 10% FCS, 1% HEPES buffer, 1% sodium pyruvate and 1% penicillin-streptomycin.

### Proteins and labelling

Rat CD2 (rCD2, residues 23-219, UniProt P08921) was cloned into the pHR plasmid with a 6H-SRAWRHPQFGG-6H tag at the C-terminus. rCD2 was expressed by lentiviral transduction using HEK 293T cells and purified using Ni-NTA beads and size-exclusion chromatography. Soluble pMHC (HLA-A) was produced as previously described ^23^. A fragment antigen binding (Fab) reactive with human CD45 was produced by digestion of Gap8.3 antibody. Both rCD2, UCHT-1 Fab and Gap8.3 Fab were labelled with Alexa 647 and 488, respectively, using the succinimidyl ester method. For bulk exclusion and calcium signaling experiments, cells were labelled with Gap 8.3 CD45 Fab at 10 μg/ml and 5 mM Fluo-4 for 15 min in phenol red-free RPMI at 37°C and subsequently washed 3 times in PBS. To perform single-molecule tracking experiments, cells were labelled with Gap 8.3 CD45 Fabs at 1 nM for 15 min and then washed 3 times in PBS.

### Imaging

#### FRAP

The mobile population in the bilayer was assessed using FRAP. Here, 10 frames were recorded prior to bleaching. Bleaching was performed by removing beam expanders. Post-bleaching, the intensity was recorded at 1 s interval over 3 min using a 100x 1.49NA Nikon TIRF objective.

#### TIRF

TIRF imaging was performed at 37°C or room temperature where indicated using 100x, 1.49NA Nikon TIRF Objective and 488 nm (Spectra-Physics-488, 100mW) and 638nm (Cobolt 06-MDL 638 nm 180 mW) lasers. The fluorescence signal was either split using a DualView2 (Photometrics) or passed through an OptoSpin spinning filter wheel (Cairn) before being recorded with a iXon Andor EMCCD camera (Andor). Image stacks were acquired at exposure times of 10 to 30 ms. When imaging with labelled Fabs, imaging was performed at room temperature within 15 min due to the high off rates.

#### Calcium imaging

Calcium imaging was performed in epi-illumination mode at 37°C using a 20x 0.5NA Plan Fluor Nikon objective with frames captured every second at 30 ms exposure for 10 min. Prior to experiments, cells were dropped onto the surfaces and imaged on OKT3 coated glass (10 min coating with 10 μg/ml OKT3, 3 times washed with PBS) to confirm the responsiveness of the cells and the fraction of cells triggering on OKT3 was generally found to be 60-80%.

### Image Analysis

#### FRAP

The mobile fraction population was calculated *via* first, correcting the recovery curve for bleaching using a reference area. The recovery curve was then normalised, taking the pre-bleach intensity (*F*_*i*_) as 1 and the intensity immediately following bleaching (*F*_*0*_) as 0. To estimate the plateau intensity (*F*_∞_) the curve was fit to a single exponential. The mobile fraction (*f*_*mobile*_) was calculated *via* ^24^

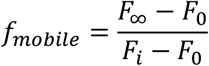

#### Particle Tracking and Diffusion Analysis

Single-particle tracking and diffusion analysis was performed using a custom-written Matlab code ^25^. Here, lipid diffusion in the bilayers was calculated by fitting the first 5 points (50 ms) of the ensemble mean-square displacement (MSD) curve to ^26^

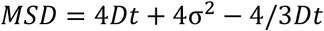

Jump-distance (JD) analysis was performed to fit 2 diffusing populations on tracks gathered from each cell. The probability distribution *P*(*r*^2^, Δ*t*) of the squared distance travelled, *r*^2^, in one time-step Δ*t* was fitted to ^25^

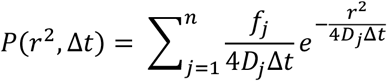

For all diffusion measurements, only tracks longer than 5 frames were used.

#### Calcium Analysis

Calcium data were analysed using custom-written software in Matlab. Here, cells were identified and tracked to gather intensity traces. Peaks where then detected using the FindPeaks in-built Matlab function. Per experiment the number of cells exhibiting a calcium response (peaks ≥ 1) and the fraction of triggering cells which exhibited multiple peaks (peaks > 1) were calculated, along with the average time to trigger the first calcium peak, integrated peak intensity and peak height in multiples of the baseline.

#### Exclusion Analysis

rCD2 and CD45 image stacks were averaged and background subtracted using the rolling background function in ImageJ (pixel size = 50). The images were then analysed using custom-written Matlab code were an Otsu ^49^ generated threshold was used to create a contact mask (rCD2 channel) and cell mask (contact + CD45 mask) and the mean intensity of CD45 inside 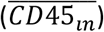 and outside of the contact 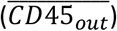 was calculated. CD45 exclusion was then set as: 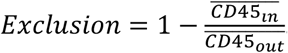.

## Data availability Statement

All raw data has been deposited on Figshare under https://figshare.com/projects/Robust_T-cell_signaling_triggered_on_soft_polydimethylsiloxane-supported_lipid-bilayers/80327

## Code availability statement

The source code for the calcium analysis has been deposited in GitHub under: https://github.com/janehumphrey/calcium.

## Supplementary information

### Supplementary videos

**Suppl. Video 1**: Lipid diffusion on 4 kPa and 1 MPa PDMS and glass supported lipid bilayers.

**Suppl. Video 2**: Diffusion of A488-tagged UCHT1 antibody Fab-coupled TCRs in rCD2 contacts formed on glass- and 4 kPa PDMS-supported lipid bilayers.

**Suppl. Video 3**: rCD2 contact formation and calcium signaling on 4 kPa PDMS-supported lipid bilayers at 37°C.

**Suppl. Video 4**: rCD2 contact formation and calcium signaling on glass-supported lipid bilayers at 37°C.

### Supplementary figures

**Suppl. Fig. 1:**
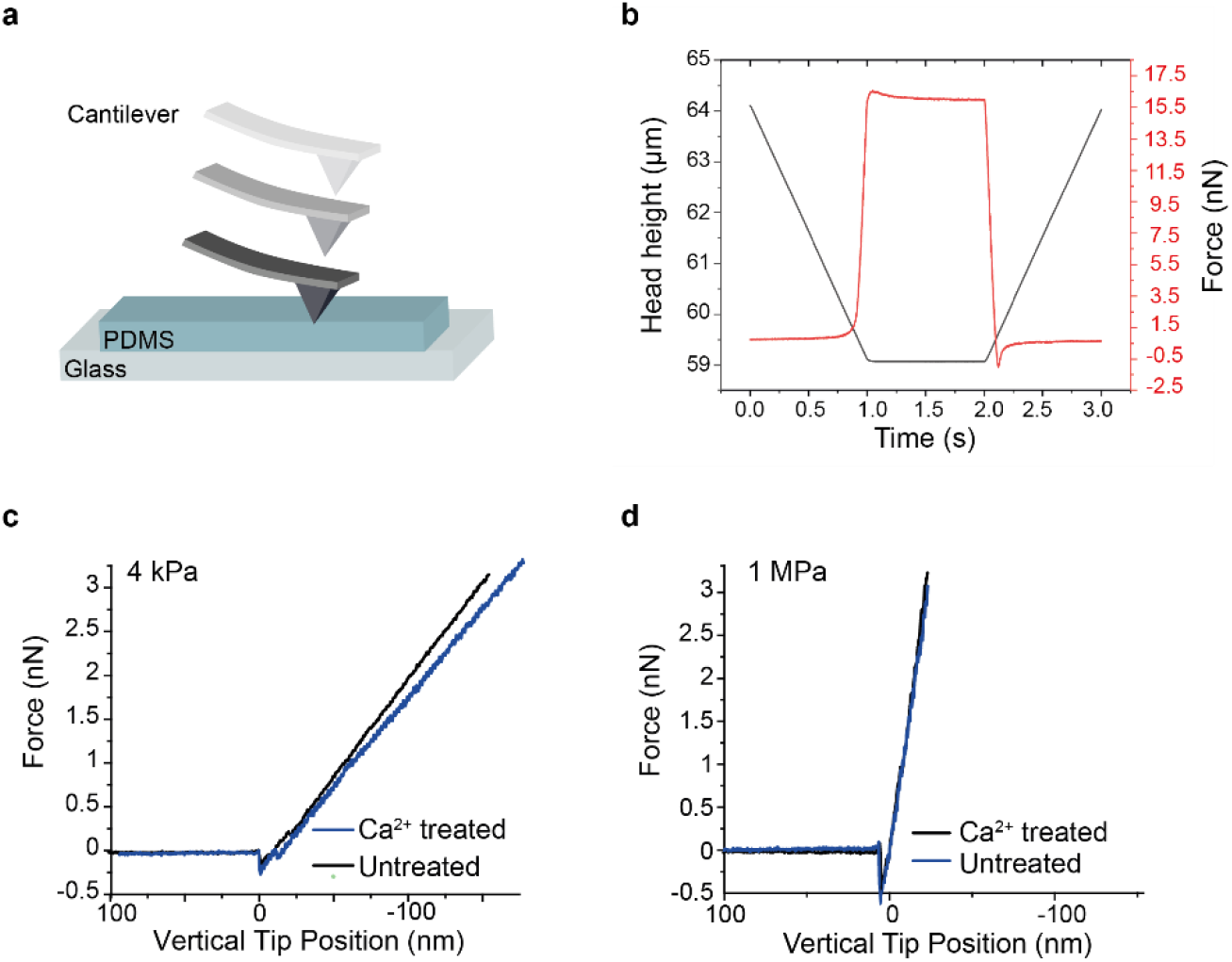
PDMS characterisation using AFM. **a** Schematic of the experiment. **b** PDMS gels exhibit a predominantly elastic response with minimal creep. **c,d** Calcium treatment of 4 kPa gel did not lead to an increase in gel stiffness for either 4 kPa or 1 MPa PDMS-supported bilayers.

**Suppl. Fig. 2:**
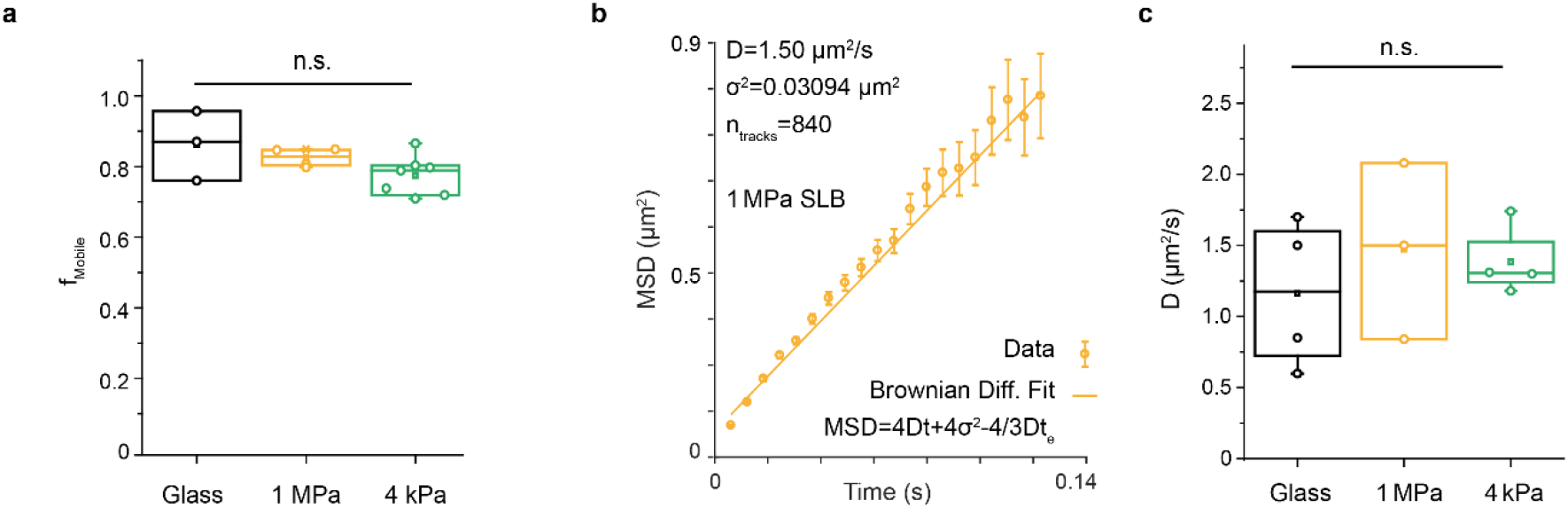
Diffusional properties of lipids are similar for glass- and 1 MPa and 4 kPa PDMS-supported bilayers. **a** The mobile population *f* assessed *via* FRAP was not significantly different between the conditions. **b** Example MSD plot of single particle tracking on a 1 MPa bilayer. **c** Diffusion coefficients gathered from single particle tracking were not significantly different between the conditions. Boxes indicate the 25% and 75% quartile, the horizontal line the mean, and whiskers the 1.5 IQR.

**Suppl. Fig. 3:**
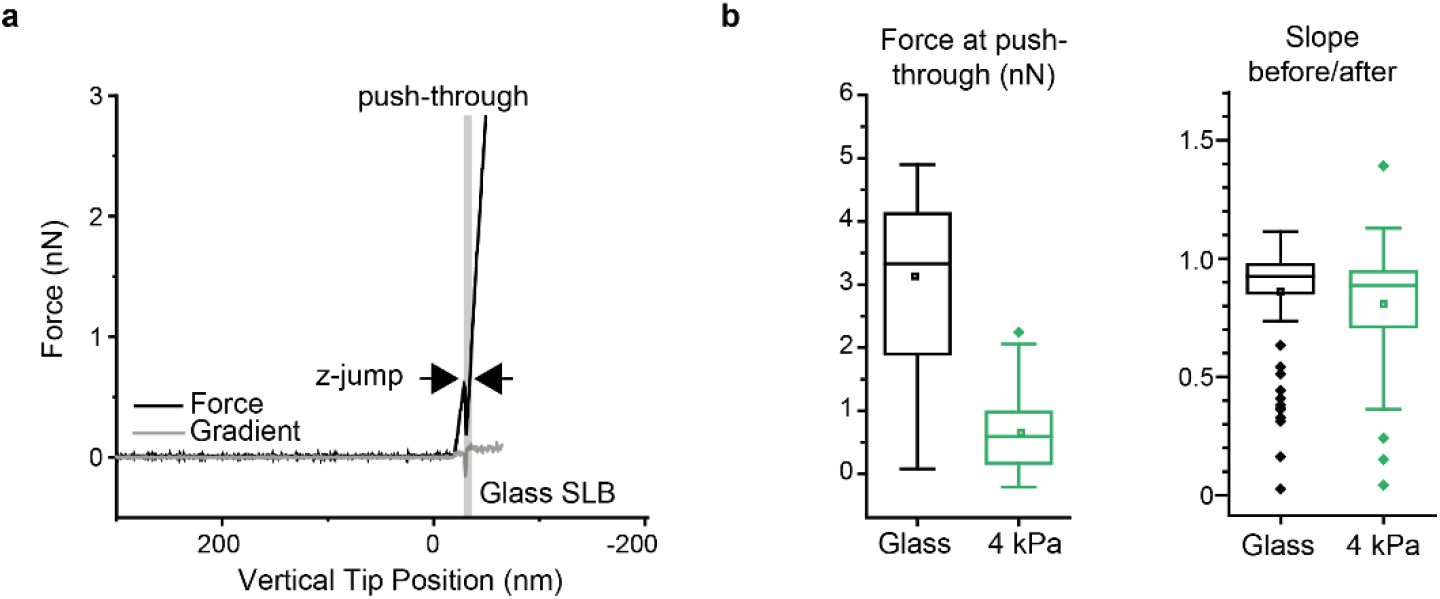
AFM-based characterisation of glass- and 4 kPa PDMS-supported bilayers. **a** Representative force trace of a cantilever pushing through a glass supported bilayer (force: black line; gradient of the force: grey). The push-through event is indicated as a “z-jump” (grey area with indicating arrows). **b,c** Boxplots comparing push-through force and slope before/after the push-through [number of force curves analysed: 52 (4 kPa PDMS), 92 (glass) from 3 independent experiments]. Boxes indicate the 25% and 75% quartile, the square is the mean, the horizontal line the median, the diamonds the outliers and whiskers the 1.5 IQR.

**Suppl. Fig. 4:**
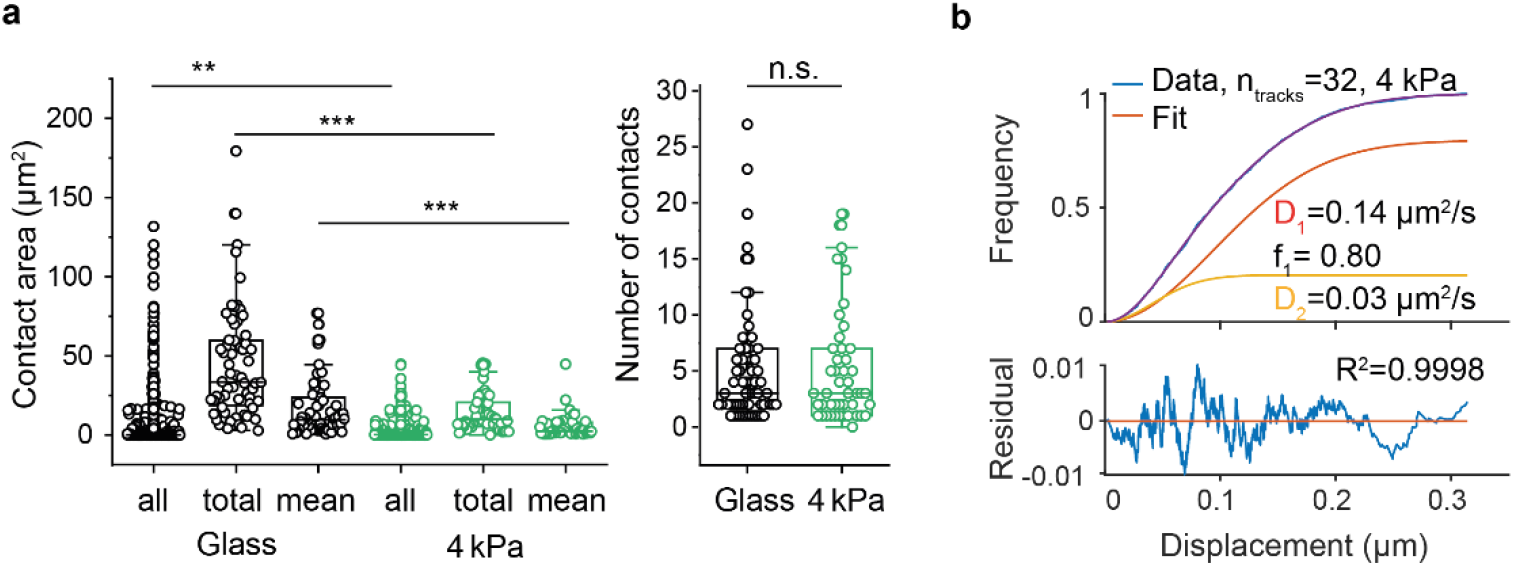
Formation of late rCD2-mediated contact areas and diffusion analysis. **a** Comparison of late rCD2 contacts: areas for all individual contacts (all), total contact area per cell (total), and average contact size (mean) were each measured (glass-supported bilayers n_cells_=64, 4 kPa PDMS-supported bilayers n_cells_=47) as well as the number of contacts per cell (right). Each data point represents one cell. Boxes indicate the 25% and 75% quartile, the horizontal line the mean, and whiskers the 1.5 IQR. The Mann-Whitney U test was used for statistical analysis with *:<0.05, **:<0.01, ***:<0.001, ***:<0.0001. **b** Example of JD analysis.

**Suppl. Fig. 5:**
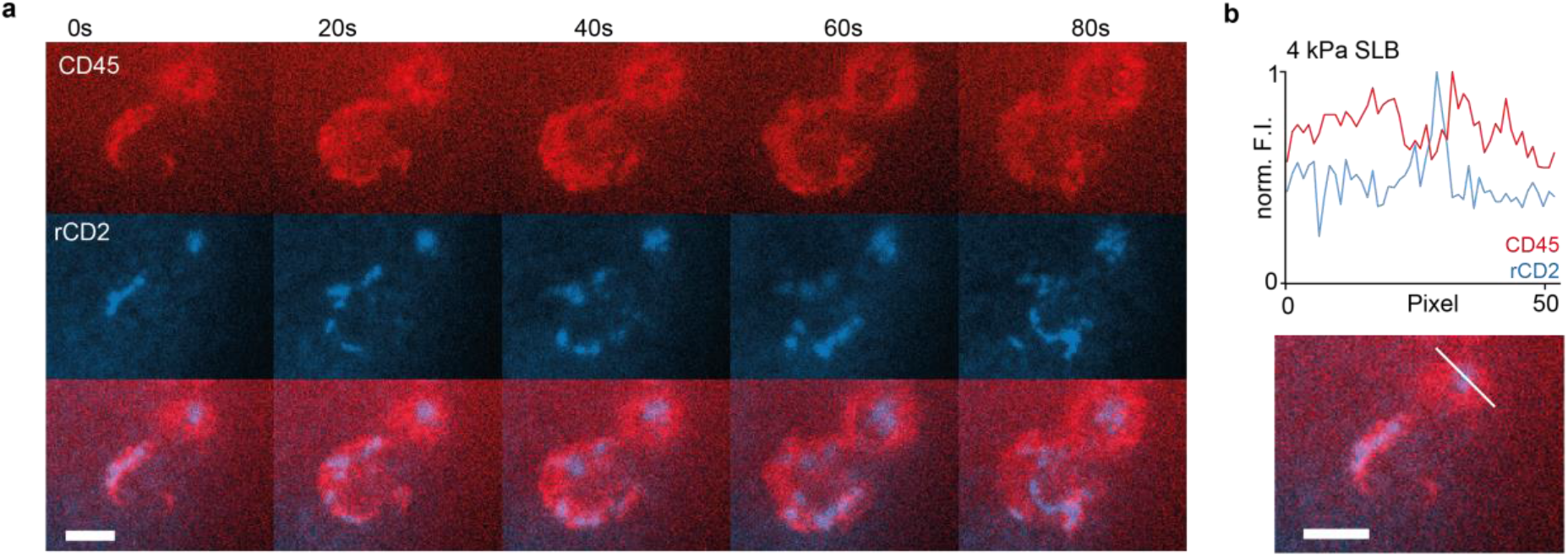
Small contacts exclude CD45. **a** Representative false-colour two-colour time-lapse TIRF imaging of a T cell recruiting rCD2-Alexa647 (blue) into contacts, alongside exclusion of CD45 labelled with Gap8.3 anti-CD45 antibody Fab tagged with Alexa 488 (red). **b** Line profile of CD45 and rCD2 fluorescence at the 0s timepoint. Scale bars 1μm.

**Suppl. Fig. 6:**
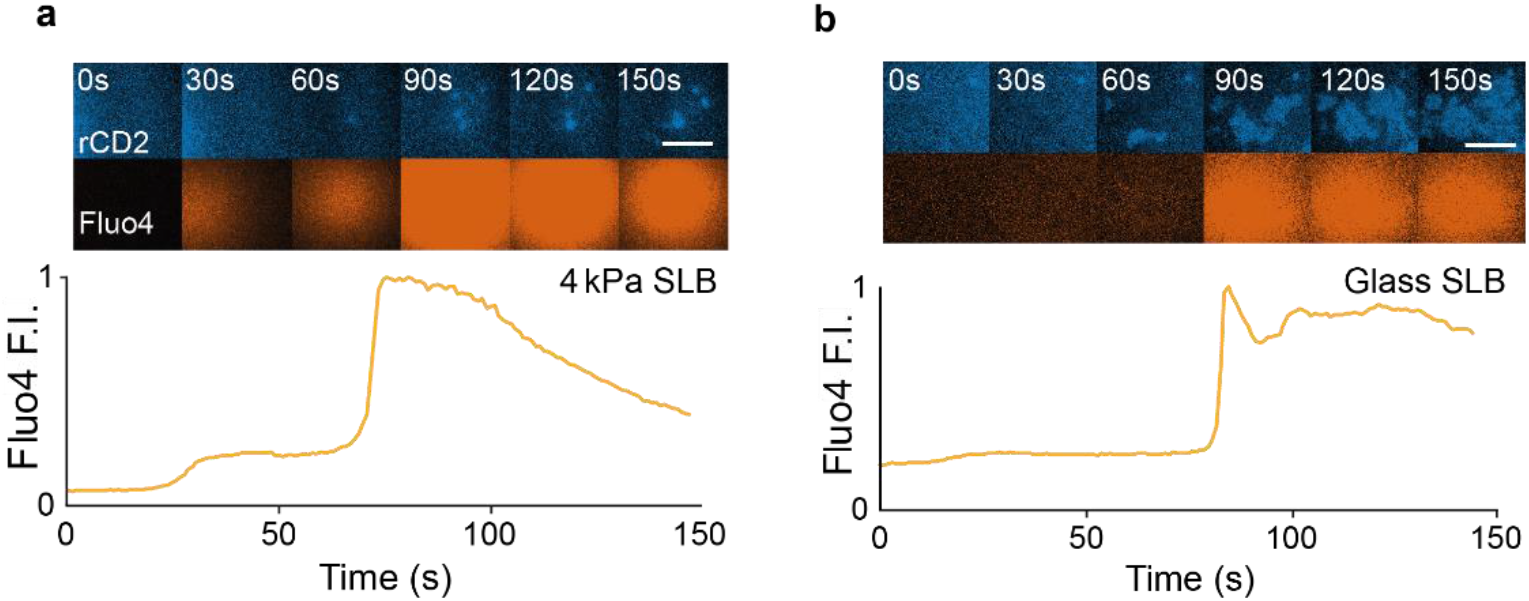
Small rCD2 contacts exclude CD45 and produce calcium responses on 4 kPa PDMS-supported bilayers. Jurkat cells expressing rCD48 were loaded with Fluo-4 and allowed to settle onto Alexa647-tagged rCD2-presenting 4 kPa PDMS (**a**) and glass (**b**) supported bilayers. Two colour illumination with a 488 laser set to epi illumination and a 647 laser set to TIRF illumination allowed simultaneous detection of rCD2 accumulation and calcium signaling. The Fluo4 signal, as raw images and traces (orange) are shown under the time-lapse TIRF images. Scale bars 5 μm.

**Suppl. Fig. 7:**
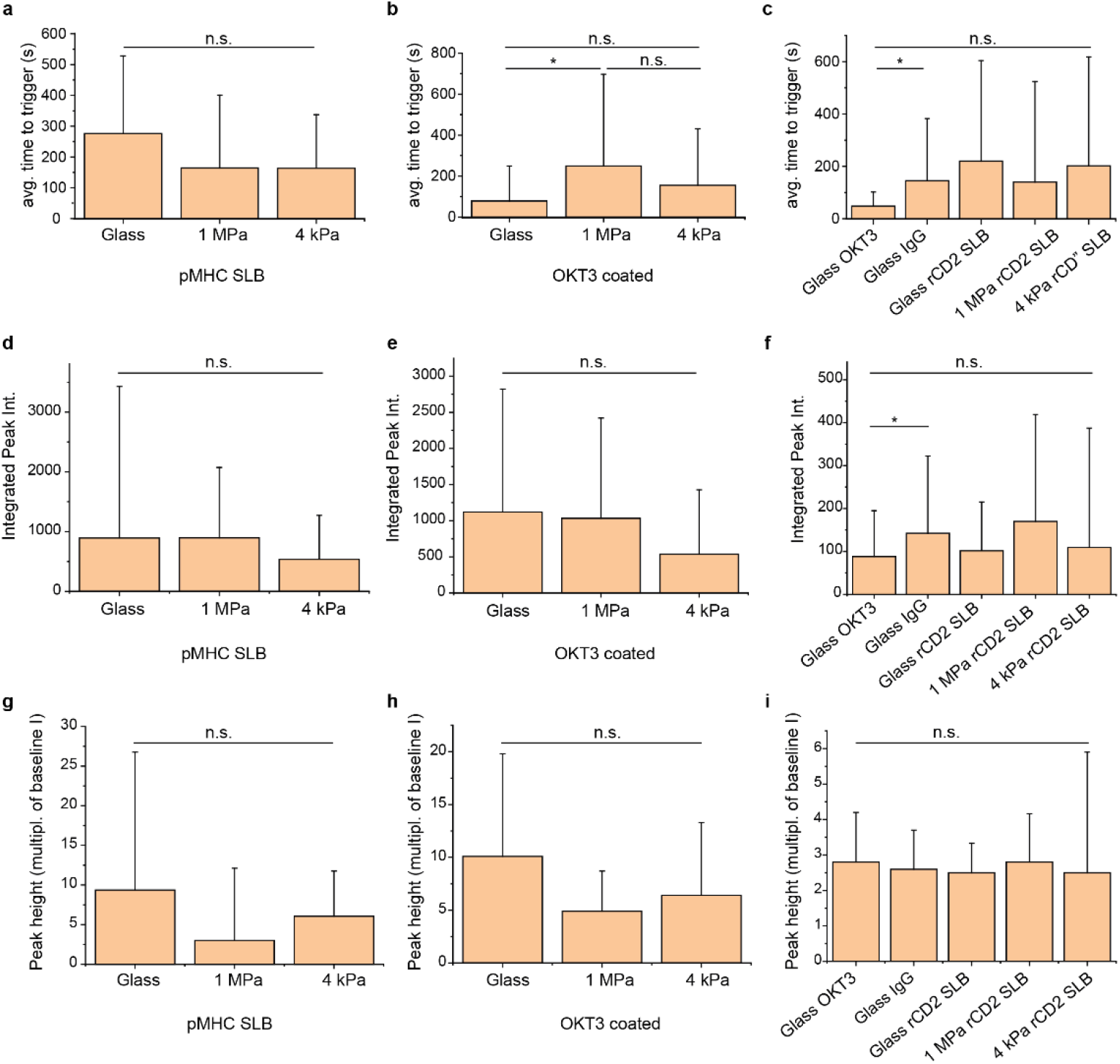
Calcium peak parameters. **a-c** Average time to trigger, **d-f** integrated peak intensity, **g-I** peak height in multiples of baseline intensity for T cells settling onto: **a,d,g** pMHC-presenting bilayers formed on glass and 1 MPa and 4 kPa PDMS; **b**,**e,h** OKT3 antibody-coated glass and 1 MPa and 4 kPa PDMS; and **c,f,I** rCD2-presenting glass and 1 MPa and 4 kPa PDMS supported bilayers as well as OKT3- and unspecific bovine IgG-coated glass. Boxes represent the average time to trigger across 3 independent experiments for n_cells_>50, with error bars representing the propagated standard deviation over the 3 experiments. All experiments were performed at 37°C. The Mann-Whitney U test was used for statistical analysis with *:<0.05.

